# The interplay of chromatin phase separation and lamina interactions in nuclear organization

**DOI:** 10.1101/2021.03.16.435657

**Authors:** Rabia Laghmach, Michele Di Pierro, Davit A Potoyan

## Abstract

The genetic material of Eukaryotes is segregated into transcriptionally active euchromatin and silent heterochromatin compartments. The spatial arrangement of chromatin compartments evolves over the course of cellular life in a process that remains poorly understood. The latest nuclear imaging experiments reveal a number of dynamical signatures of chromatin which are reminiscent of active multi-phase liquids. This includes the observations of viscoelastic response, coherent motions, Ostwald ripening and coalescence of chromatin compartments. There is also growing evidence that liquid-liquid phase separation of protein and nucleic acid components is the underlying mechanism for the liquid behavior of chromatin. In order to dissect the organizational and dynamical implications of chromatin’s liquid behavior, we have devised a phenomenological field-theoretic model of nucleus as a multi-phase condensate of liquid chromatin types. Employing the liquid chromatin model of *Drosophila* nucleus, we have carried out an extensive set of simulations with an objective to shed light on the dynamics and chromatin patterning observed in the latest nuclear imaging experiments. Specifically, we demonstrate that the emergence of the chromatin mesoscale channels and spheroidal droplets could arise from the interplay of chromatin type to type interactions and intermingling of chromosomal territories. We also shed light on the dynamical heterogeneity and coherent motions of chromatin domains which are captured by micro-phase separation of chromatin types. Finally, we further illuminate the role of heterochromatin-lamina interactions in the nuclear organization by showing that heterochromatin-lamina interactions enhance the mobility of euchromatin and indirectly introduce correlated motions of heterochromatin droplets.

## I. INTRODUCTION

The nuclei of eukaryotic organisms have hierarchically organized genomes, the structure and dynamics of which correlate with the phenotypes and cellular functions [1, 2]. At the top of the hierarchical order, the nuclear space is partitioned into chromosome territories within which one finds deeper layers of polymeric chromatin order, including micro-compartments, nano-scale sub-compartments, and loops [3–5]. At the nanometer resolution, chromatin is seen as an amorphous and dynamic organization of poly-nucleosomal arrays, the nature of which is poorly understood [6, 7]. At micron scales, however, chromatin shows a relatively simpler picture which is commonly rationalized in terms of two distinct mesoscopic states of chromatin [8]: a diffuse genetically active euchromatin and dense genetically inactive heterochromatin.

The existence of a discrete number of mesoscale chromatin states is further supported by the Hi-C experiments [9] which reveal 1D patterns of type A and type B chromosomal loci defined by distinct epigenetic tags. Polymer simulations using the A/B type decorated co-polymeric sequences have shown that 1D sequence of A/B types contain sufficient information for recapitulating both the 3D chromosomal folds and the dynamics at the euchromatin/heterochromatin borders [10, 11]. It is known that concentrations of euchromatin (EC) vs. heterochromatin (HC) in the nuclei are generally time and cell line dependent [12]. The heterochromatin is known to phase separate into facultative (fHC) and constitutive (cHC) forms with former typically residing in the nuclear periphery because of attractive interactions with nuclear lamins while the later is localized in chromocenters [13]. This spatial organization of euchromatin/heterochromatin regions is what is known as the conventional nuclear architecture. A notable exception to the conventional architecture has been found in the nuclei of nocturnal mammals which display inverted architectures [12, 14, 15] with heterochromatin concentrating at the core of the nucleus. Given that the nuclear organization is not random one naturally anticipates a physical connection between euchromatin/heterochromatin distribution and its temporal dynamics. The super-resolution imaging techniques such as Hi-D [16], reveal a heterogeneous variation of chromatin organization with dynamics which appears to be more consistent with a picture of a multi-phase liquid condensate [17–20]. The reports of ubiquitous condensations of proteins and nucleic acids in the nucleus through liquid-liquid phase separation (LLPS) [21–25] leave little doubt that multiple aspects of nuclear structure and function are dictated by the liquid behavior and multivalent interactions of nuclear proteins and RNA with the chromatin [26–30].

The notion of liquid chromatin states [27] is further supported by the observations of the viscoelastic response of chromosomal loci [11, 31–33], coherent motions of chromatin domains [32, 34], the coalescence and Ostwald ripening of chromatin droplets [35, 36], the existence of epigenetic zonations and chains of interlinked ∼ 200 − 300 nm wide chromatin domains reminiscent of polymer melts [20]. The presence of non-equilibrium, motorized ATP-driven processes is also being reported to modulate chromatin dynamics which manifests in the form of ATP-dependent flows, driven fluctuations and anomalous diffusion coefficients of chromatin loci [34, 37].

To dissect the role of the chromatin’s liquid behavior on the global order and dynamics of nuclear chromatin domains in this work, we develop a field-theoretic description of the nucleus as a liquid condensate of chromatin types. By carrying out simulations with multiphase liquid chromatin model of nucleus, we elucidate a number of recently observed features in nuclear imaging experiments. Namely, we show (i) conditions favoring the segregation of heterochromatin droplets vs. connected mesoscale chain states of chromatin domains [12, 20] (ii) the role of fluctuations and heterogeneous diffusive dynamics of chromatin loci [34, 37, 38] (iii) emergence of coherent motions across nuclear compartments [32]. (iv) the impact of heterochromatin-lamina preferential positioning on intranuclear chromatin order [39].

## II. A PHASE FIELD THEORY FOR LIQUID CHROMATIN CONDENSATES

In this section, we describe the physical foundation and the mathematical formulation of the field-theoretic model of nucleus as a multi-phase condensate of chromatin types. The present model is partially based on the previous mesoscale formulation of nucleus proposed by us [40] which has introduced a single liquid chromatin type for studying chromatin boundary fluctuations and territorial compartmentalization during growth and senescence of *Drosophila* nucleus. In the present formulation, the nucleus is resolved at a chromatin subtype level corresponding to facultative/constitutive heterochromatin and euchromatin forms. The primary driving forces for emergent nuclear architecture and dynamics are derived from microphase separation of heterochromatin sub-types, surface tension of chromatin droplets, and differential affinity for chromatin-lamina interactions. Volume and surface constraints are imposed on chromatin types for capturing chromosomal and nuclear boundaries. Given the dense, active and heterogeneous nature of nuclear chromatin, it is worth highlighting the advantage of field-theoretic description which manages to avoid the notorious glassy states encountered in the particle based polymer simulations, thereby facilitating the study of long-timescale chromatin dynamics and patterning at the scale of whole nucleus [41, 42].

The field theoretic description of nucleus in the present work is best described in terms of experimentally motivated constraint on shapes and sizes of nucleus and chromosomes which are supplemented by the physically motivated interaction terms accounting for polymer intermingling and movement within and between chromosomal territories. To define the shape of the nucleus, we introduce an auxiliary order parameter *η* which takes 0 inside the nucleus and 1 outside, and varies smoothly between these two values through the interfacial region. This region represents the nuclear envelope whose position is given by the iso-contour *η* = 1*/*2. The field *η* is used as an indicator function independent of time to model a fixed nucleus with volume *V*_*N*_. Since the nuclear envelope is assumed here at the equilibrium state during interphase of the cell cycle, it can be represented by a tanh-like profile that corresponds to the analytical shape of a diffuse planar interface that separates the two bulk phases 0 and 1. Thereby, we simulated the oblate nuclear shape typical of eukaryotic nuclei during interphase by the use of the following expression: 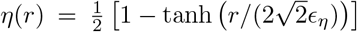; Where 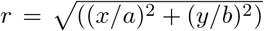 is the distance from the center of the physical domain Ω, whereas *a* and *b* are the semi-major and semi-minor axes. The width of the nuclear envelope is given by the constant 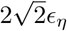.

Within the nucleus, the N-chromosome territories are described by an N-dimensional vector of non-conserved order parameters ***φ***(r, *t*) = {{*φ*_*i*_} _*i*=1,*…,N*_}. The microphase separation of A, B and C chromatin types within chromosome territories is resolved through two additional non-conserved order parameters ***ψ*** (**r**, *t*) = {{*ψ*_*j*_} _*j*=1,2_} which quantify the epigenetic state of the chromosomal region. Similarly, the phase-field variables *φ*_*i*_(r, *t*), *i* = 1, …, *N*, and *ψ*_*j*_(**r**, *t*), *j* = 1, 2, vary smoothly across their interfaces profile between two values that it is 1 inside its domain and 0 elsewhere.

The dynamics of the chromatin compartmentalization patterns is derived from the free energy functional ℱ which describes the intra-nuclear phase separation of chromatin subtypes. The constructed free energy functional for nuclear chromatin is an extension of our previous model [40], and can be split into two free energy functionals *F*_*B*_ and *F*_*I*_. The Ginzburg-Landau free energy functional *F*_*B*_ describes the coexistence of two phases associated to each phase-field variables completed by volume constraints terms to ensure the shapes change is given by:

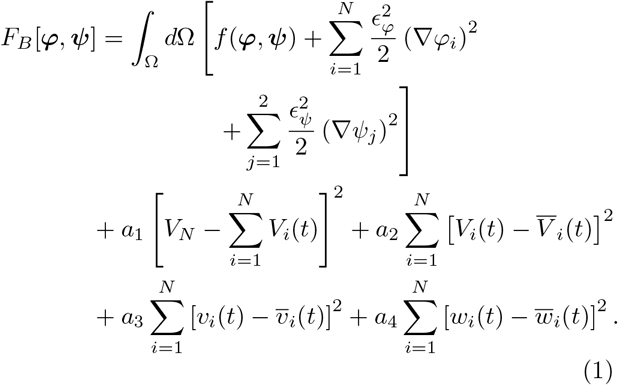

where *f* (***φ, ψ***) is the bulk free energy contribution for multi-phase field variables, and gradients terms accounting for the presence of different interfaces in the system and contributing to the interfacial energies. The gradient parameters *E*_*φ*_ and *E*_*ψ*_ are controlling the thickness of the interface profile of ***φ*** and ***ψ***, respectively. For the bulk free energy density, we use a multi-well potential expressed as: 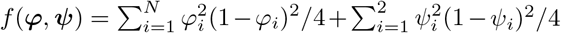. The terms proportional to *a*_*i*_ account for volume constraints required to enforce the volume of the chromosomal territories at their prescribed values 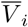, the facultative heterochromatin at 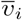, and the constitutive heterochromatin at 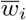. The parameters *a*_1_, *a*_2_, *a*_3_ and *a*_4_ are positive coefficients that control the thermodynamic driving forces of coarsening processes of different compartments present in the nucleus. The volumes of the *i*-chromosomal territory *V*_*i*_(*t*), facultative and constitutive heterochromatin domains within each chromosome *v*_*i*_(*t*) and *w*_*i*_(*t*) are defined as the spatial integral over the physical domain Ω of their interface profiles given by the associated phase-field variables *φ*_*i*_(**r**, *t*), *ψ*_1_(**r**, *t*) and *ψ*_2_(**r**, *t*). Using the usual phase-field approximation of the volume could change the coexistence phases values defined by the phase-field variables in 0 and 1. We thus used an interpolation function, defined as 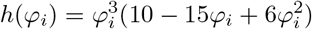, for approximating the volumes of different compartments of the nucleus while keeping the position of the local free energy minima at the coexistence phase values. Thus, the volumes of these domains are approximated by the following expressions: *V*_*i*_(*t*) = ∫ _Ω_ *d Ω h*(*φ* _*i*_); *v*_*i*_(*t*)=∫ _Ω_ *d Ω h*(*φ*_*1*_); *h*(*φ*_*i*_); and *w*_*i*_(*t*) = ∫ _Ω_ *d Ω h*(*φ*_*2*_); *h*(*φ*_*i*_).

Next we define the free energy functional *F*_*I*_ which accounts for the geometrical constraints on the nucleus, excluded volume interactions between different domains within the nucleus, and the interaction between heterochromatic subtypes with the nuclear envelope. The functional *F*_*I*_ is expressed as:

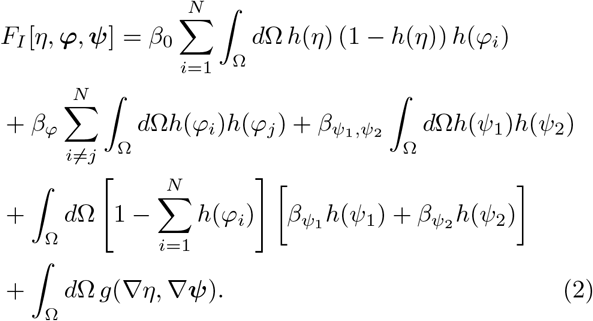

The first term in *F*_*I*_ corresponds to the energy penalty reflecting a geometrical constraint on the nuclear volume which is required to restraint nuclear components’ motion inside the nucleus. The other terms represent the excluded volume interactions between chromosome territories, fHC-cHC regions interactions and HC/EC regions mixing affinity, with the interactions strengths described by the parameters 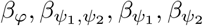, respectively. The last term represents the interaction of the fHC and cHC with the nuclear envelope though a *g*-function. This function represents the local Lamina-interaction energy contribution to the free energy functional, and it is expressed as:

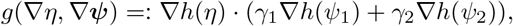

where *γ*_1_ and *γ*_2_ are two positive parameters controlling the binding affinity of heterochromatin types to the nuclear lamina.

The full free energy functional of the nuclear chromatin ℱ [*η*, ***φ, ψ***] that we minimize to get the evolution equations of the nuclear structures is equal to the sum of the two energy functional contributions *F*_*B*_ and *F*_*I*_ explicitly presented in Eqs. (1−2). The phase-field parameter *η* that describes nuclear geometry is assumed constant in the total free energy functional. The dynamics of nuclear chromatin is governed by the Allen-Cahn evolution equations of the phase-field variables {***φ, ψ***}:

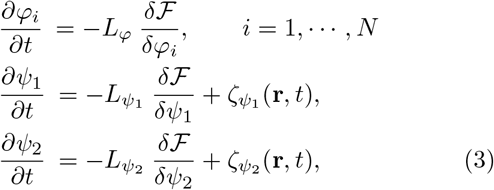

where 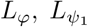 and 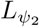 are mobility coefficients which are proportional to the relaxation time of different phase-field variables. We have chosen the mobility coefficients to match the magnitudes of *in vivo* measurements of euchromatin and heterochromatin diffusion coefficients [43] The terms 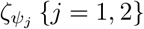 in Eq. (3) account for the fluctuations at the boundaries of EC/fHC and EC/cHC islands due to finite size nature of the droplets. The fluctuations are modeled as white noise: 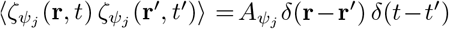 where the amplitude of noise is given by: 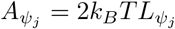. the amplitude of noise *A*_*p*_ sets the “effective temperature ” *T*_*eff*_ of the nucleus [44] which can be taken as a measure of ATP activity in comparisons with the experiment [45, 46].

### Parametrization of the *Drosophila* nucleus

To solve numerically the set of evolution equations of the phase-field variables resulting from the free energy minimization (3), we use the finite element method combined with the preconditioned Jacobian Free Newton Krylov technique (JFNK). The model has been implemented in MOOSE finite element C++ library [47, 48] which is built using high-performing computational libraries MPI, LibMesh, and PETSc needed for solving non-linear partial differential equations [48]. In this work, all simulations were performed on a rectangular mesh domain of dimension 6 *µm* × 9 *µm*. The same domain mesh as the one in [40] is used here: a quadrilateral elements are chosen to generate the fine mesh with 640000 elements. The time-step used for time integration is set to 0.04 in dimensionless units which is chosen to insure numerical stability of all simulations. The nuclear shape is maintained in an elliptical shape in which the semi-major axes are fixed to *a* = 2.5 *µm* and the semi-minor axis to *b* = 4 *µm*; thus the nuclear volume is: *V*_*N*_ = *πab* = 31.416 *µm*^2^ consistent with empirical measurements of *Drosophila* nucleus during interphase [49].

For simplicity, the mobility of chromosomal and heterochromatin compartments are set to be equal 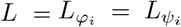. This parameter fixes the domain’s interface relaxation time: *τ* ∝ *L*^*−*1^. The interface relaxation time *τ* is used to set the unit timescale for the kinetics of phase transitions. To investigate effects of diffusion coefficient of chromatin on nuclear morphology, given by 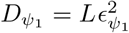 for the heterochromatin and 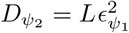 for chromocenter, simulation were performed by varying the interfacial parameters 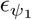 and 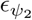. The diffusion coefficient values taken for A/B types of heterochromatin are 4, 12, 20 *µm*^2^*/s*. The diffusion coefficient of chromosomes *D*_*φ*_ is fixed to 1 *µm*^2^*/s*. These values of diffusion coefficients are motivated by experimental measurements of euchromatin/heterochromatin mobility in live cells [43]. The interaction coefficient between chromosomal territories and nuclear envelope is chosen to be strong enough to maintain them inside the nucleus by setting *β*_0_ = 66.7. To ensure a well separated chromosome territories, we set the chromosome-chromosome interaction parameter with strong interaction same as the one in [40] *β*_*φ*_ = 40. In previous work [40], we have shown that strong affinity between euchromatin-heterochromatin leads to mixed states for heterochromatin droplets while weaker mixing affinity leads to their fusion. We thus set the heterochromatin-heterochromatin interaction parameter 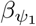 to 0.1 for all simulations, and chromo-chromocenter and hetero-chromocenter interaction parameters 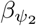 and 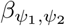 are varied. The interaction parameters between A/B types of heterochromatin and nuclear lamina, *γ*_1_ and *γ*_2_, are varied to evaluate the effects of competition between binding energies and chromatin compartment interactions. The remaining model’s parameters including the growing domain kinetic are set as *a*_1_ = 0.16 and *a*_2_ = *a*_3_ = *a*_4_ = 2. The fluctuation amplitude of euchromatin-heterochromatin *A*_1_ is fixed at 15 and two differnt values are considered for euchromatin-chromocenter interface *A*_2_ to 2.5 and 15. We note that while the nuclear shape, size, chromosome numbers and diffusion coefficients are calibrated after *Drosophila* nucleus the model is sufficiently general for drawing broader conclusions about the chromatin structure and dynamics in eukaryotic nuclei.

## III. RESULTS

### A. Chromatin compartmentalization patterns emerge from phase separation of heterochromatin types

To dissect the impact of the phase separation of heterochromatin types on spatial organization of chromatin in the nucleus, we first carry out simulations by varying the free energy terms controlling the interaction of cHC-cHC, fHC-fHC, and cHC-fHC in the absence of lamina-heterochromatin anchoring terms. The interactions between chromosome territories (CTs) and the thickness of chromatin intermingling regions have a signficant impact on the interchromosomal interactions by coordinating chromatin domain formation. Therefore, we first dissect the contribution of each of these interactions on the chromatin organization. Subsequently we will use the distinct classes of nuclear architectures as stepping stone for more detailed investigation of phase separation kinetics, heterohromatin patterning and the role of lamina-heterochromatin interactions in the later sections. We note that the the concept of chromosome territories and the degree of chromatin intermingling in the current multiphase liquid chromatin model is essentially controlled through a single parameter *ϵ*_*φ*_.

The evolution of nuclear chromatin patterns driven by the attractive interactions between the two heterochromatin types cHC and fHC is shown in Fig. 2A and with repulsive interaction in Fig. 2B. The simulated nuclear patterns are generated for two extreme cases of cHC mixing affinity which is set by the parameter 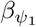. We note that the mixing affinity parameters 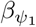 and 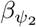 represent the degree of the attractive interaction between different heterochromatin regions from neighborhood chromosomes. The simulation results with a stronger and weaker mixing affinity are shown in the top and the down panels of the figure 2 A–B), respectively. Depending on the strength of chromosomal territorial interaction *β*_*φ*_, the decrease of mixing affinity parameters leads to enhanced mobility of heterochromatin droplets to move through chromosome territories. Thus, it will be easy for similar-type heterochromatin from different chromosomes to merge and initiate the phase separation. This connection between similar-type heterochromatin from neighboring chromosomes is shown in the figure 2. We observe that for an attractive interaction between cHC-fHC, with a stronger mixing affinity 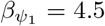, the fHC droplets located initially on neighbouring chromosome centers move toward periphery of their territories and coalesce forming a larger droplet. Meanwhile the cHC droplets remain in the center (Fig. 2A top panel). For a similar mixing affinity of heterochromatin types, which are set to ensure a weaker mixing within a chromosome 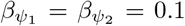, both of the heterochromatin droplets are moving through the chromosomal periphery in the direction of the nuclear center resulting in a single droplet (Fig. 2A bottom panel). In contrast, a strong interaction between cHC and fHC heterochromatin (Fig. 2B) leads to phase-separated states of chromatin compartments irrespective of the mixing affinity of heterochromatin types. One can also observe an increase in the distance between heterochromatin droplets when the cHC-fHC interaction is made stronger.

**FIG 1.**
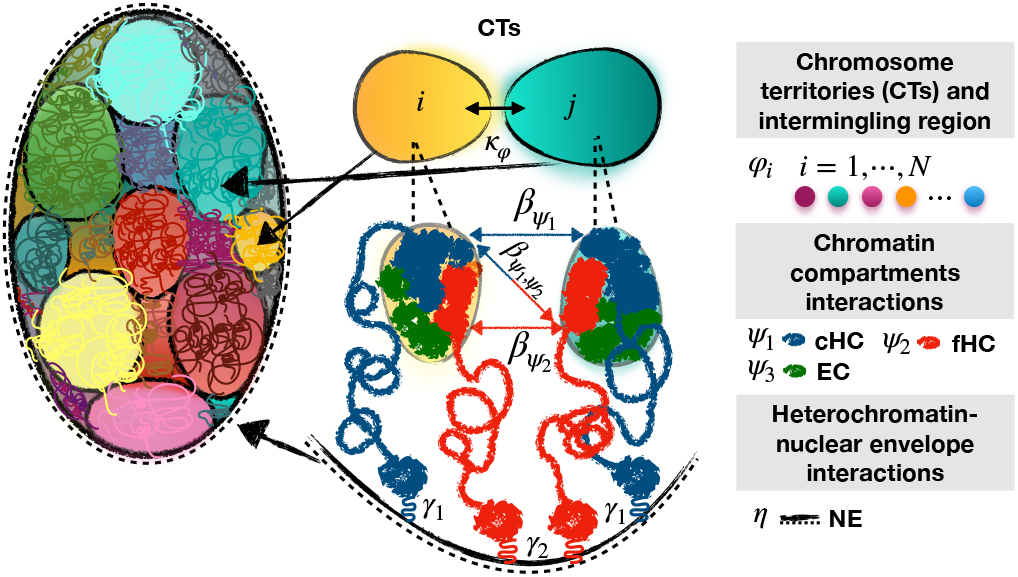
Schematic representation of the mulitphase liquid chromatin model of nucleus. Shown are the key physical interactions that make up the global nuclear free energy functional. Namely, the chromosome territorial interactions (CT), interactions between constitutive and facultative heterochromatin types (cHC-cHC, cHC-fHC and fHC-fHC) and the lamina interaction which is modeled via surface gradient terms modulated by *γ*_1_(cHC-lamina) and *γ*_2_(fHC-lamina).

**FIG 2.**
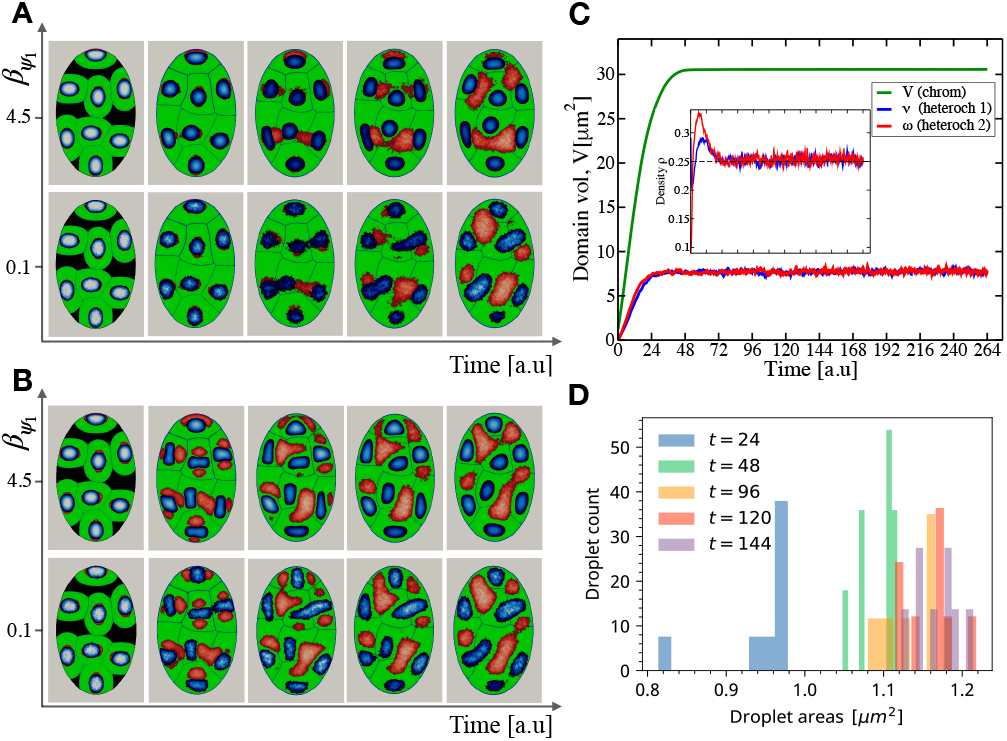
Compartmentalization patterns of nuclear chromatin. The snapshots show the emergence of chromatin compartmentalization at different time steps initiated with attractive and repulsive interactions between the cHC and fHC heterochromatin types. A) Simulated structures obtained for attractive interaction between cHC-fHC with a weaker mixing affinity of cHC (top panel) and stronger mixing affinity (bottom panel) B) Simulated structures obtained with repulsive interaction between cHC-fHC with a weaker mixing affinity of cHC heterochromatin (top panel) and stronger mixing affinity of cHC (bottom panel). The green color indicates the chromosome territories, while the blue and red colors indicate the heterochromatin types cHC and fHC regions, respectively. C) Relaxation dynamics of chromosomal volume *V*, the cHC volume *v* and fHC volume *w* D) Hetero-chromatin droplet heterogeneity quantified as area distribution as a function of simulated time.

In order to fix the total time for simulating interphase nuclear dynamics as well as to assess the temporal convergence compartmentalization patterns, we have also analyzed the temporal evolution of the chromosome territories volume *V*, the fHC and cHC heterochromatin volumes *v* and *w* respectively (Fig. 2C) and the distribution of fHC droplet areas (Fig. 2D). The different volumes of the nucleus’ components are relaxed to reach their prescribed values and thus occupy the whole volume of the cell nucleus. Likewise, the relative fraction of the two heterochromatin types whose we have evaluated by *v/V* and *w/V* are well around 25%.

### B. Thermodynamics of chromatin phase separation dictates coherent motions of chromatin domains

Here we study the role of differential mobility of chromatin types on the kinetics of nuclear compartmentalization. To vary the mobility we first express the diffusion coefficients of chromatin types in terms of mobility coefficients 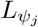 which allows fixing both the relaxation times of heterochromatin fields *ψ*_*j*_ and the diffusive interface thickness *ϵ*_*j*_, such that 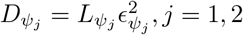. With this formulation, we can vary these two parameters independently for setting the diffusion coefficients. Simulations were performed with different values of diffusion coefficients associated with the cHC and fHC heterochromatin domains by varying their diffusive interface thickness 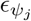. In these simulations, we consider only the case of repulsive cHC-fHC interactions with an identical mixing affinity for cHC and fHC types.

We introduce and use local displacement vector fields of chromatin to analyze the chromatin domains’ motion within the nucleus and the resulting compartmentalization patterns. The local displacement fields of chromatin domains are correlated to their interface’s motion; therefore, they can be determined by the interface’s normal velocity. We use the following expression to characterize the velocity fields of heterochromatin droplets [50, 51]:

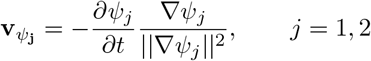

The velocity fields of heterochromatin domains are proportional to the diffusion coefficients via the term *∂ψ*_*j*_*/∂t*. The figure 3 shows that quite naturally the heterochromatin droplets move faster to the nucleus’s center for an increased diffusion coefficient values. As can be seen, a decrease in the number of identical-types of heterochromatin droplets with increased values of diffusion coefficient, which reveals that the diffusion accelerates the kinetics of heterochromatin droplet coalescence. Despite the self-attractive interaction of fHC domains (weak mixing affinity of fHC and cHC), we find that the fHC-droplets are fully disconnected in all of the chromosome territories when one has a large diffusion coefficient of cHC relative to fHC (5 times or higher). The cHC-droplets, on the other hand move rapidly to the nuclear center and coalesce.

**FIG 3.**
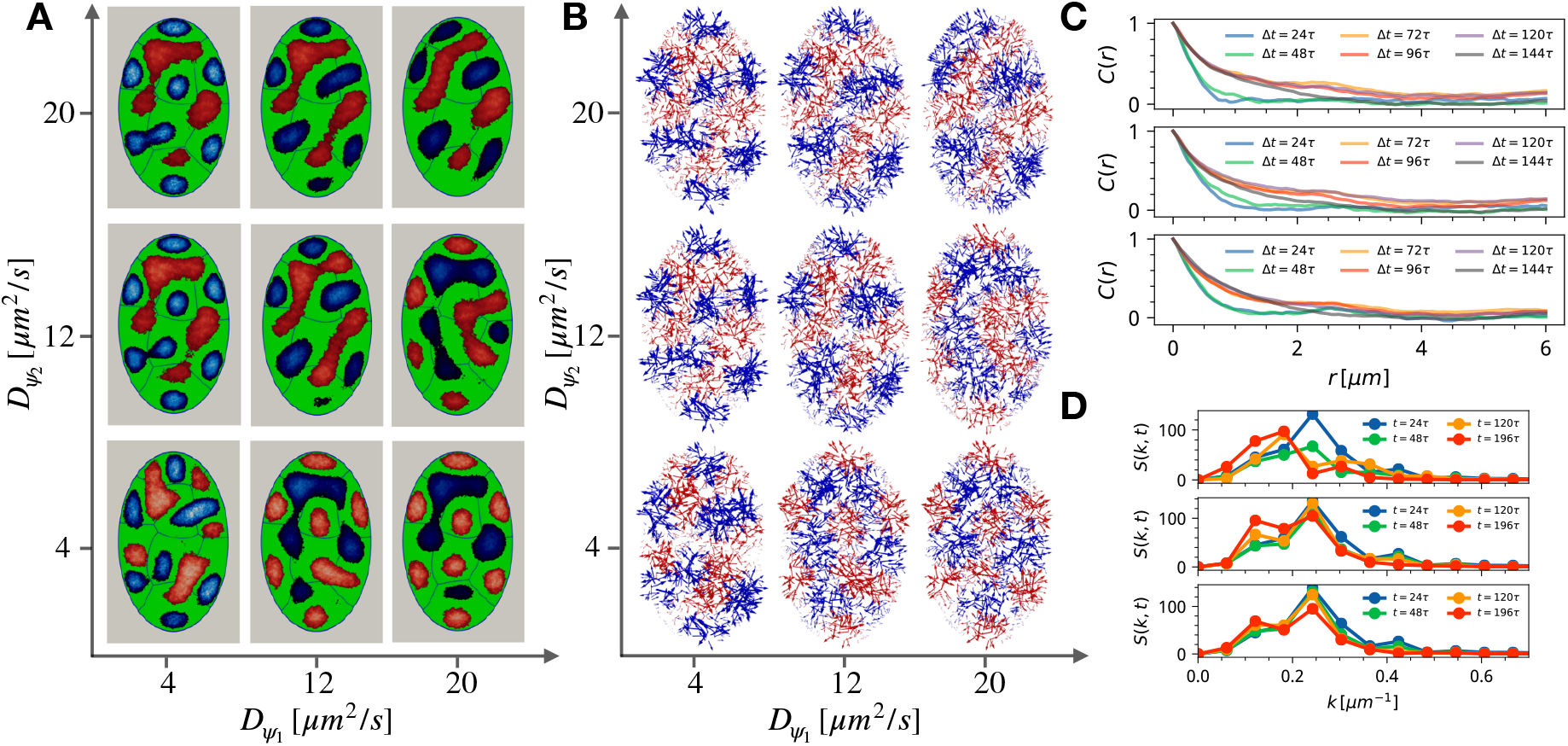
The role of heterochromatin diffusion and fluctuations on the emergent subnuclear chromatin morphology and dynamics. A) Shown are nuclear chromatin patterns for repulsive interaction between cHC and fHC heterochromatin types and weaker mixing affinity of both cHC and fHC. (B) Displacement fields of cHC and fHC corresponding to one-time unit. Shown are simulation results from fast intermediate and slow diffusion coefficients corresponding to panels arranged from top to bottom, respectively. (C) Correlation functions of displacement fields as a function of the displacement period. (D) Dynamic structure factors of heterochromatin domains. Shown are results from fast intermediate and slow diffusion coefficients corresponding to panels arranged from top to bottom, respectively.

We calculate the spatial correlation functions for heterochromatin domains 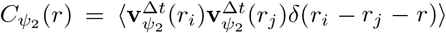 by performing ensemble averaging over the velocity vectors 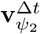 defined over the physical domain Ω for different displacement periods Δ*t*. The spatial auto-correlation functions for heterochromatin (either cHC or fHC) capture the experimentally observed growth and eventual decline of correlations over the nuclear scale [32]. The growth of correlations has previously been shown in polymeric simulation with their origin clearly stemming from the microphase separation of chromatin types [11]. However, the polymeric simulations have shown growth of correlation length with no decline which is likely an artifact due to finite size of systems and glassy dynamics which often plagues simulations of dense polymer systems in confinement. Herein we are able to confirm that indeed the microphase separation of chromatin domains is responsible at least in part in generating correlation motions of domains and the emergence of mesoscopic Dynamically Associated Domains (DADs) [11] which do indeed decline over micron scales of nucleus. The impact of chromatin mobility seems to damp the smaller time-scale correlations with the overall long time-scale trends dictated by phase-separation kinetics. The most conspicuous role of the diffusion coefficient is revealed by analyzing dynamical structure factors *S*(*k, t*) which show that chromatin domains with higher mobility favor the formation of connected mesoscopic channels over a fixed period of time (Fig 3D).

### C. The differential mobility and lamina interactions of heterochromatin types contribute to conventional nuclear order

The crucial role of nuclear lamina and lamin-heterochomatin interactions in controlling nuclear chromatin architecuture is widely acknowledged [14, 15]. Here we examine the impact of interactions of distinct forms of heterochromatin with lamina located at the nucelar envelpe (NE) on the global nuclear chromatin patterning. To this end we have carried out simulations by varying the affinity coefficients *γ*_1_ and *γ*_2_ controlling the tethering of cHC and fHC domains to the NE, respectively. In the following simulations, we consider all the same interaction parameters of chromatin domains used in the previous sections. We note that the cHC is modeled at a higher effective temperature with respect to fHC set by the amplitude of flucutatons. Hence the model is capable and does break the symmetry of nuclear architectures over finite times for the swapping parameters of heterochromatin-lamina interactions.

The nuclear morphology resulting from the phase separation, which is initiated by the chromatin type to type interactions, and the interactions with the nuclear lamina are shown in figure 4A. The resulting morphology illustrates how the interaction strengths of different heterochromatin compartments with the NE affect the chromatin positioning relative to the nuclear periphery. We see that quite naturally the distinct types of heterochromatin, cHC and fHC, move closer to the nuclear periphery and tether to it due to the attractive interactions (*γ*_*i*_ *>* 0). The contact layer between heterochromatin regions and the nuclear envelope tends to increase with the binding affinity coefficients. We quantify accumulation of heterochromatin at nuclear periphery via radial profiles (4B) which are computed as the azimuthaly averaged histograms of heterochromatin types measured from the center of nucleus obtained from the final simulation mesh configuration: *P* (*r*) = *N*_*i*_(*r*)*/M* (*r*) where *N*_*i*_(*r*) is the pixel count of heterochromatin type i (cHC or fHC) which is at a distance [*r, r* + Δ*r*] from the center of nucleus and *M* (*r*) is the total pixel count of the mesh at at a distance [*r, r* + Δ*r*].

**FIG 4.**
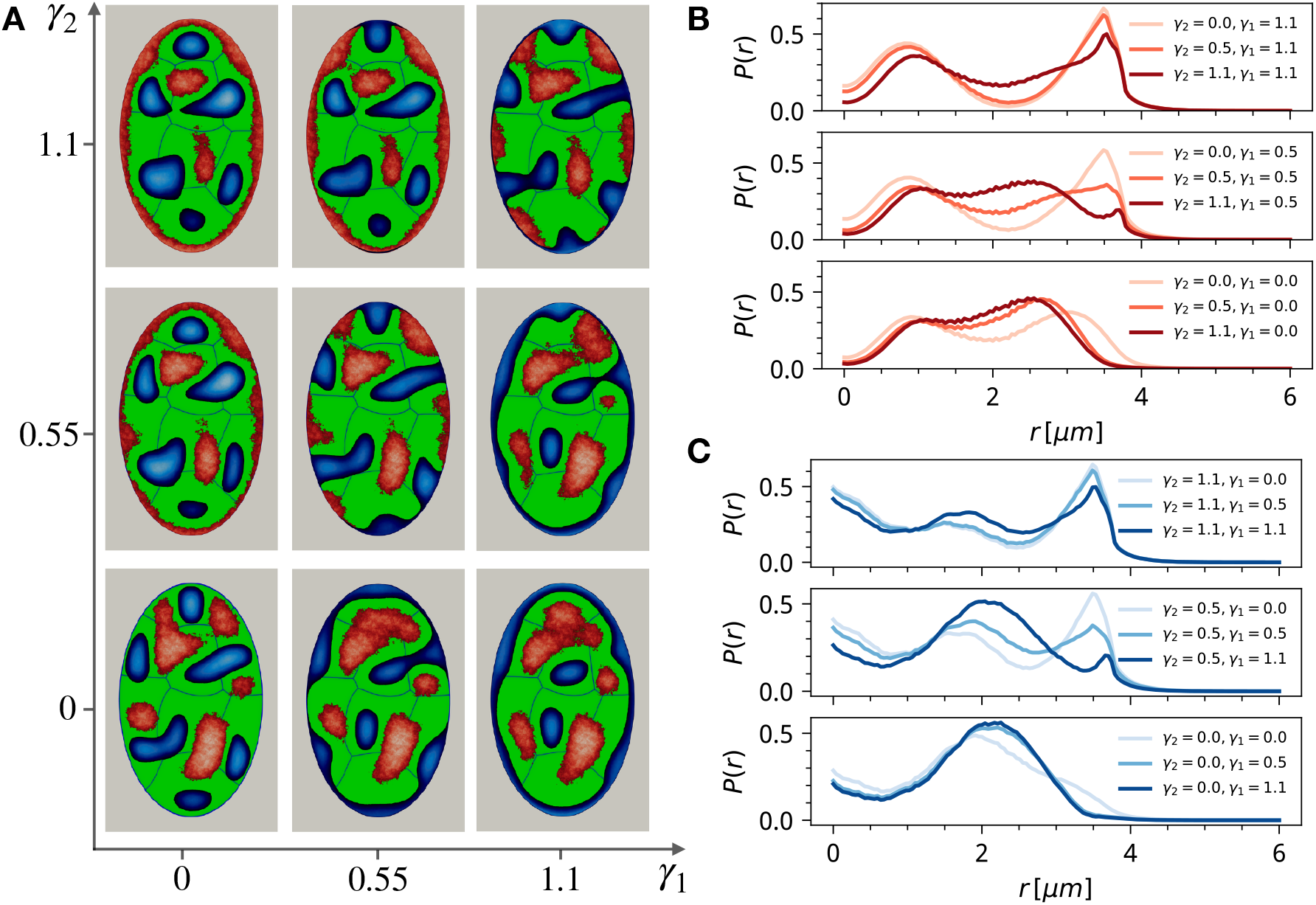
The role of lamina-heterochromatin interactions on nuclear chromatin patterns. (A) Shown are nuclear morphologies generated for various strengths of nuclear lamina interactions with cHC domain *γ*_1_and fHC domain *γ*_2_. (B) Radially averaged density profile of cHC measured from a center of nucleus. (C) Radially averaged density profile of fHC measured from a center of nucleus.

Turning on the attractive interaction of cHC type with the lamina *γ*_1_ *>* 0 in the absence of fHC lamina interaction *γ*_2_ = 0, facilitates the formation of nuclear patterns where cHC is localized at the periphery of NE with a thickness of layer depending on the strength of interaction parameter *γ*_1_. Since *γ*_2_ = 0, the fHC droplets move through their chromosomal territory in the direction of the nuclear center and coalesce with neighboring droplets forming a larger droplet. The cHC droplets are experiencing motion towards the NE because of the attractive interaction with the NE. However, the cHC droplets located initially in the nucleus center have less freedom to move to NE due to the strong interaction with nearest fCH droplets. Consequenstly, the fCH droplets far from the NE remain at the center of their chromosome territory. We obtain a similar patterning when turning on interaction of fHC types with lamina while turning off the same interaction for cHC *γ*_1_ = 0 and *γ*_2_ *>* 0. When an equal strength of NE attractive interaction is considered i.e. *γ*_1_ = *γ*_2_, we find that the cHC and fHC droplets attach to the NE with alternating positions of the droplets. Furthermore, depending on the strength of NE attractive interaction with the cHC and fHC droplets, we can simulate a predominant attachment of a given heterochromatin type to the nuclear lamina. For instance the fHC droplets are attached to almost the whole NE’s surface and arranged as a layer near it when *γ*_2_ *> γ*_1_. However, only the cHC droplets very close to the NE attach to it in this case, since they interact relatively weakly with the attractive NE compared with fHC droplets. From the simulations with varying lamina-heterochromatin interactions, we learn that the interplay between the chromatin type to type interactions located in different chromosomal territories and heterochromatin domain interactions with the nuclear lamina have a significant impact the global nuclear patterning (Fig. 4). Indeed, the competition between these interactions determines how droplets position relative to the NE as well as the thickness of heterochmatin layers attached to NE.

### D. Heterochromatin content controls the percolation threshold in the connectivity of chromatin droplets

The fractional content of chromatin types in the nucleus is expected to strongly impact the nuclear organization. Here we focus specifically on the connectivity of droplets and the ability to form mesoscale chromatin channels [12, 20] which are natural candidates for gene regulation through chromatin architecture. To investigate the role of heterochromatin fraction in the nuclear architecture, we performed simulations by varying the total heterochromatin fraction which includes the constitutive and facultative components for two limiting strengths of the cHC-fHC type interactions. The Lamina interactions are set to *γ*_2_ = 2 × *γ*_1_ = 1.1 which allow tethering of heterochromatin components to the NE. The self-interactions of cHC and fHC domains are set to the same values used in previous section. The simulated nuclear architectures are presented in figure 5 A). The corresponding profiles of the phase-field variables *ψ*_1_ and *ψ*_2_ along the major-axis of the nucleus are shown in figure 5 B).

**FIG 5.**
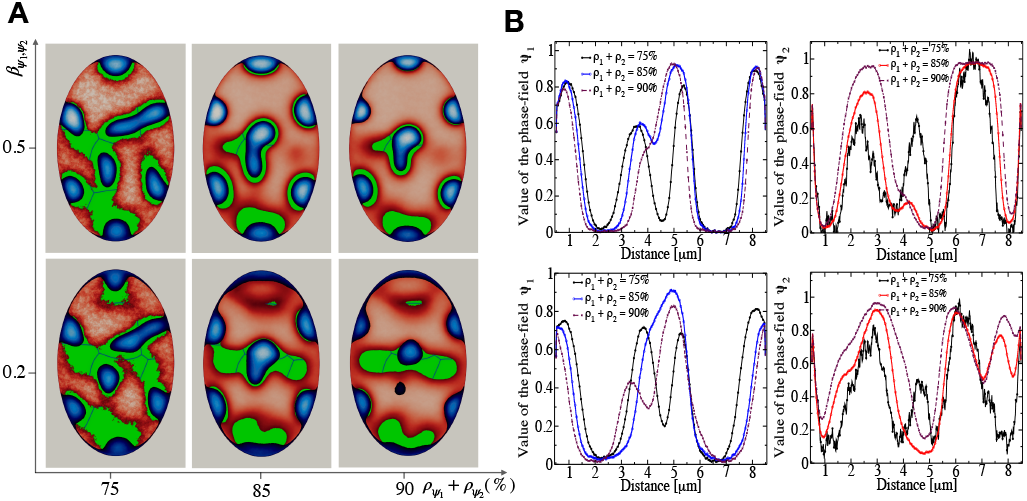
Variation of the average density of cHC and fHC heterochromatin types. A) Shown are simulated nuclear chromatin patterns for different values of heterochromatin content fraction in the nucleus as a function of interactions strength between cHC-fHC domains. B) Profiles of the phase-field variables *ψ*_1_and *ψ*_2_along the major-axis of the nucleus as a function of heterochromatin density. The profile plots are associated with the simulated morphology presented in A) in the same order as the strength of the cHC-fHC interactions.

The resulting morphology shows a non-linear dependence on the relative volumes occupied by cHC and fHC domains and the cHC-fHC interaction parameter 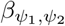 (Fig. 5 B). For weak interaction between the cHC and fHC, we find that the cHC droplets are stretched along the NE forming a thin connected layer. The connectedness and thickness of the layer is a function of fHC content. We find that for large fractional contents there is a well percolation threshold for the dominant form of heterochromatin. This percolation threshold is modulated by the cHC-fHC. Specifically the strong interaction between the cHC and fHC prevents the fusion of cHC droplets by limiting their freedom to move along the NE. From simulation results, we can conclude that both the lamina interactions and the heterochromatin density contained in the nucleus can have a significant consequence in how intranuclear droplets are arranged and how much they will be dynamically constrained during the interphase cycle of the cell.

## IV. CONCLUSION

The 4D organization of chromatin in eukaryotic nucleus shapes the gene expression and phenotypes states of cells. Several independent observations suggest that on the micrometer scales chromatin exhibits behaviour reminiscent of a multi-phase liquid condensates. The liquid phases in the nucleus arise from the distinct epigenetic tags on the chromatin. The chromatin types are a defining feature of cellular phases, yet their connection with the organization and temporal evolution of nuclear architectures remains poorly understood.

In the present contribution, we have developed a phenomenological field-theoretic model of nucleus which resolves chromatin types as liquid states and therefore allows us to dissect the role of chromatin liquid behavior on the emergent nuclear architectures and dynamics. With the multiphase liquid model of nuclear chromatin, we are able to explicitly track motions and interactions of chromatin types both within and outside chromosome territories as well as with the nuclear lamina.

An extensive set of simulations with multi-phase liquid chromatin model of *Drosophila* nucleus reveals that an interplay between the affinity for phase separation and the intermingling of chromosomal domains can give rise to a wide range of architectural patterns observed in interphase eukaryotic nuclei. Specifically, we are able to recapitulate patterns with segregating heterochromatin droplets as well as mesoscopic channels recently observed in the super-resolution nuclear imaging experiments [20].

Besides generating distinct nuclear architectures, we also show that the model readily captures many dynamical observables which emerge from the interplay of phase separation and interactions of heterochromatin types with the lamina. Remarkably, the model is able to capture the full spectrum of time scales for the experimentally observed coherent motion of heterochromatin domains which displays an initial growth phase followed by an eventual decay of spatial auto-correlation functions [32].

The simulations have also shed light on the contribution of heterogeneous heterochromatin-lamina interactions in interphase nuclear architecture. We show that heterochromatin-lamina interactions, besides the more obvious consequence of generating spatial gradients of heterochromatin, also have a less obvious effect of raising the effective mobility of euchromatin which facilitates micro-phase separation and correlated motions of unbound heterochromatin domains.

In summary, our multi-phase liquid model of nucleus has shown promise at capturing some of the more salient features of emerging nuclear order and dynamics. This motivates to pursue a line of quantitative refinements where one could, for instance, begin to account for the observed non-equilibrium motorized forces due to nuclear actin, inter-conversion between heterochromatin types as well as spatially dependent droplet viscosity in order to establish a more direct link with chromatin phase density and hydrodynamics.

## ACKNOWLEDGMENTS

Research reported in this publication was supported by the National Institute Of General Medical Sciences of the National Institutes of Health under Award Number R35GM138243. This work used the Extreme Science and Engineering Discovery Environment (XSEDE), which is supported by the National Science Foundation grant number ACI-1548562 [52] on the Stampede2 machine at the Texas Advanced Computing Center (TACC) through allocation CTS190023. The authors also acknowledge financial support from Iowa State University.

## Notes

### Competing Interest Statement

The authors have declared no competing interest.

